# Deep Learning-powered Bessel-beam Multi-parametric Photoacoustic Microscopy

**DOI:** 10.1101/2021.12.21.473705

**Authors:** Yifeng Zhou, Naidi Sun, Song Hu

**Affiliations:** Department of Biomedical Engineering, McKelvey School of Engineering, Washington University in St. Louis, MO 63130 USA

**Keywords:** Bessel beam, deep learning, multi-parametric photoacoustic microscopy

## Abstract

Enabling simultaneous and high-resolution quantification of the total concentration of hemoglobin (C_Hb_), oxygen saturation of hemoglobin (sO_2_), and cerebral blood flow (CBF), multi-parametric photoacoustic microscopy (PAM) has emerged as a promising tool for functional and metabolic imaging of the live mouse brain. However, due to the limited depth of focus imposed by the Gaussian-beam excitation, the quantitative measurements become inaccurate when the imaging object is out of focus. To address this problem, we have developed a hardware-software combined approach by integrating Bessel-beam excitation and conditional generative adversarial network (cGAN)-based deep learning. Side-by-side comparison of the new cGAN-powered Bessel-beam multi-parametric PAM against the conventional Gaussian-beam multi-parametric PAM shows that the new system enables high-resolution, quantitative imaging of C_Hb_, sO_2_, and CBF over a depth range of ∼600 μm in the live mouse brain, with errors 13–58 times lower than those of the conventional system. Better fulfilling the rigid requirement of light focusing for accurate hemodynamic measurements, the deep learning-powered Bessel-beam multi-parametric PAM may find applications in large-field functional recording across the uneven brain surface and beyond (*e.g*., tumor imaging).

## I. Introduction

Over the past decades, photoacoustic microscopy (PAM) has been proven to be a promising imaging modality for biomedical research due to its ability to provide functional and physiological imaging *in vivo* at the microscopic level [1]–[4]. Recent developments of multi-parametric PAM have enabled simultaneous quantification of the total concentration of hemoglobin (C_Hb_), oxygen saturation of hemoglobin (sO_2_), blood flow, and the associated tissue oxygen metabolism [5]– [7]—carving out a unique technical niche for PAM.

In the multi-parametric PAM, the microscopic resolution is typically achieved by focusing a Gaussian-shaped laser beam into a micron-level spot, where a tradeoff between the lateral resolution and the depth of focus exists due to light diffraction. As a result, when the region of interest is out of focus, the resolution and signal-to-noise ratio (SNR) degrade rapidly. Moreover, since the quantitative measurements of blood oxygenation and hemodynamics relies heavily on the resolution and SNR [8], they become inaccurate in the out-of-focus region, placing a practical limit on the application of multi-parametric PAM.

The out-of-focus issue can be addressed by performing two-dimensional raster scanning at multiple depths (*i*.*e*., z-stack imaging). However, the image acquisition time increases proportionally, making it inapplicable to studies requiring high imaging speed. An electrically tunable lens can be used to speed up this process by quickly shifting the light focus axially [9]; however, its response time is not sufficient to follow the MHz laser repetition rate. Alternatively, three-dimensional contour scanning has been developed to address the out-of-focus issue caused by uneven tissue surfaces [8], [10], [11]. However, this approach requires the addition of a motorized linear stage along the vertical axis and the development of sophisticated signal processing for online detection of the surface contour and real-time feedback to the motorized linear stage. More importantly, it cannot address the out-of-focus issue due to the limited depth of focus associated with the Gaussian-beam excitation. On the other hand, Bessel-beam excitation has been widely utilized to extend the depth of focus in intravital light microscopy [12], [13], and was recently adapted for PAM [14]. However, the previous studies were focused on structural imaging, leaving its utility in functional and quantitative imaging unexplored. In addition to the Bessel-beam excitation, deep learning has also been used to address the out-of-focus issue associated with the Gaussian beam [15], [16]. In particular, conditional generative adversarial networks (cGAN) have been successfully applied to biomedical image analysis [17]–[19]. The integration of Bessel-beam excitation and cGAN offers a promising solution to the enhancement of depth of focus.

In this article, we report a cGAN and Bessel-beam excitation-based multi-parametric PAM. First, *in vivo* data concurrently acquired by Gaussian- and Bessel-beam excitation were used to train a cGAN adapted from Isola *et al*. [20] for suppressing the side lobes and enhancing the depth of focus of the Bessel beam. Second, a side-by-side comparison of the multi-parametric PAM systems using Gaussian- and Bessel-beam excitation was carried out to examine if the Bessel-beam multi-parametric PAM was able to achieve accurate functional imaging over an extended depth range, besides the known benefit in structural imaging. Finally, cortex-wide imaging was performed over the live mouse brain to demonstrate the benefit of cGAN-powered Bessel-beam multi-parametric PAM in applications where large-field functional recording across the uneven brain surface is needed.

## II. Methods

The architecture of the cGAN is shown in Fig. 1, which is adapted from the original model [20]. The B-scans excited by Bessel-beam at multiple depths are used as the input, and the B-scans excited by Gaussian-beam in the focal plane are used as the target. In our case, the image sizes of both the input and output are set to 256 × 256 × 1. Specifically, for the *in vivo* application, the number of images for training is 26,800. The number of epochs is set to 10. The architecture of upsample and downsample layers remains the same as in the original model. The random jitter function is disabled to avoid scaling and cropping because each pixel in the image has a fixed step size. For the total generator loss, the weight of the mean absolute error between the generated image and the target image is set to 2000. Fig. 1(c) and Fig. 1(d) show the normalized photoacoustic (PA) amplitudes of training datasets in the focal plane acquired by Bessel and Gaussian-beam, respectively. After training, the cGAN is supposed to learn from the Bessel-beam excited B-scans and restore in-focus B-scans with minimal side lobes and high accuracy.

**Fig. 1.**
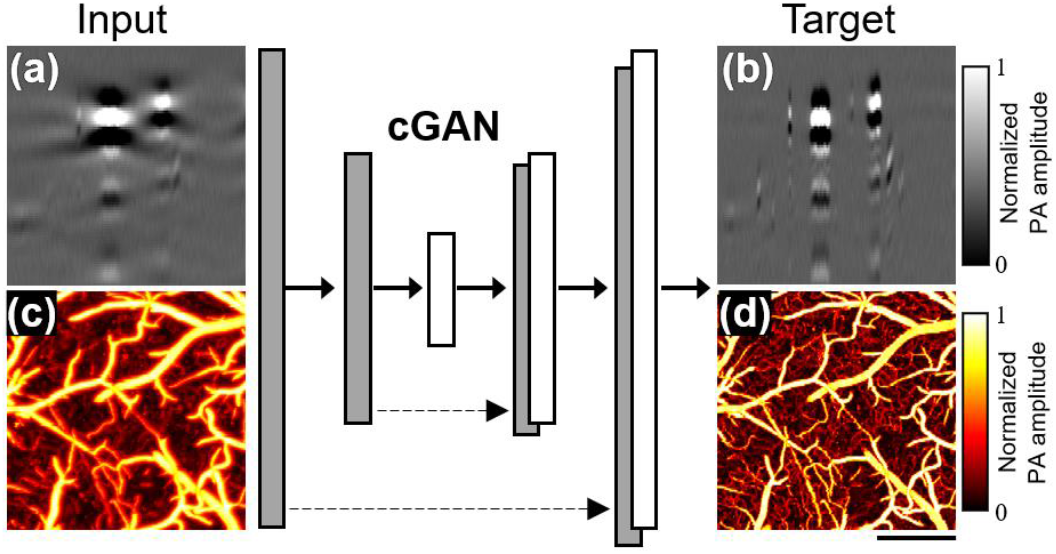
Architecture of the cGAN. The B-scans acquired by Bessel-beam excitation at multiple depths are used as the input, (a) shows one example of the B-scans in the focal plane. The B-scans by Gaussian-beam excitation in the focal plane are used as the target, (b) shows the B-scan corresponding to (a). (c) and (d) show the normalized photoacoustic (PA) amplitudes of training datasets in the focal plane acquired by Bessel and Gaussian-beam, respectively. Scale bar: 500 μm.

Figure 2 shows the schematic of our experimental setup. The Bessel-beam multi-parametric PAM utilizes a nanosecond-pulsed laser (GLPM-10-Y13, IPG Photonics; wavelength: 532 nm). Individual pulses coming out of this laser are switched between two optical paths by an acousto-optic modulator (AOM; AOMO 3080-122, Crystal Technology). When the AOM is off, the laser light passes it without diffraction (*i*.*e*., 0^th^ order) and is coupled into a polarization-maintaining single-mode optical fiber (PM-SMF, HB450-SC, Fibercore) through a fiber coupler (CFC-11X-A, Thorlabs). The stimulated Raman scattering (SRS) in the PM-SMF red-shifts the laser wavelength from 532 nm to 558 nm [21], [22]. Then, a bandpass filter (CT560/10bp, Chroma) is used to isolate the 558-nm component. When the AOM is on, ∼60% of the 532-nm light will be diffracted (*i*.*e*., 1^st^ order) into the second optical path, where no wavelength conversion is implemented. Due to the low energy, the ∼40% undiffracted light does not undergo the SRS-based wavelength conversion and thus is rejected by the bandpass filter. Thus, pulse-by-pulse switching of the laser wavelength can be realized by alternating the AOM. The two optical paths are merged by using a dichroic mirror (DM; FF538-FDi01, Semrock), and the dual-wavelength beam is coupled into a PM-SMF to maintain the linear polarization.

**Fig. 2.**
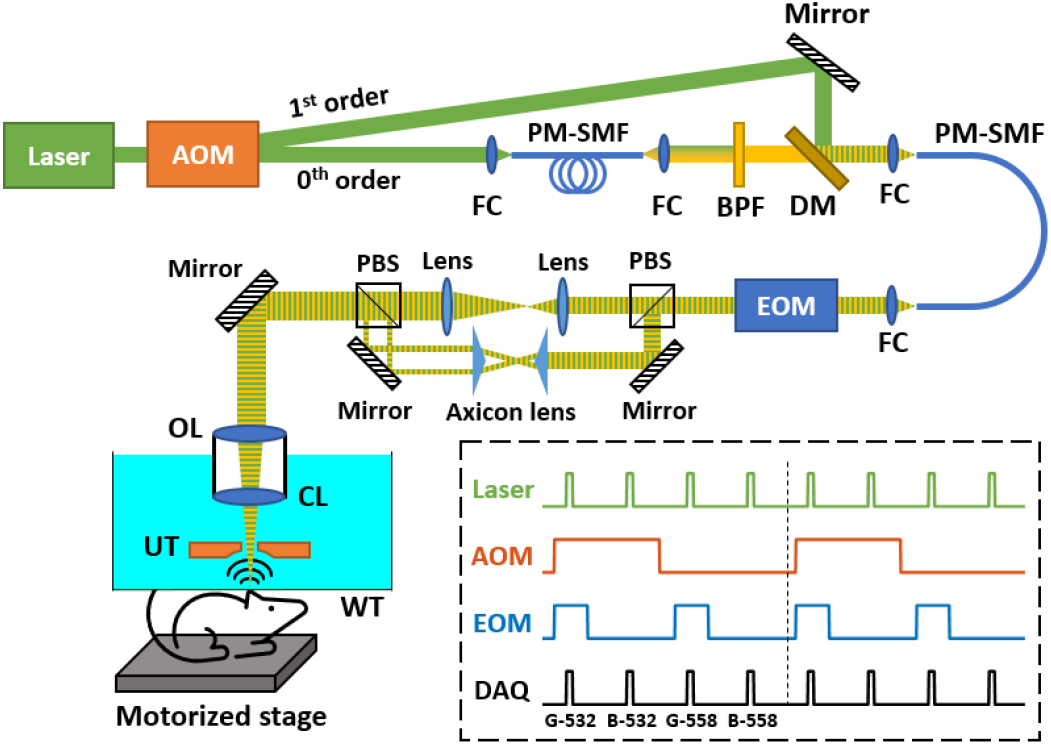
Schematic of the Bessel-beam multi-parametric PAM system. The boxed inset illustrates the laser excitation scheme designed for simultaneous Gaussian- and Bessel-beam multi-parametric PAM. AOM, acousto-optic modulator; FC, fiber coupler; PM-SMF, polarization-maintaining single mode fiber; BPF, bandpass filter; DM, dichroic mirror; EOM, electro-optic modulator; PBS: polarizing beamsplitter; OL, objective lens; CL, correction lens; UT, ultrasonic transducer; WT, water tank. DAQ, data acquisition.

After collimation of the dual-wavelength beam, an electro-optic modulator (EOM, model 350-80, Conoptics) is used to rotate the polarization. When a low voltage (*i*.*e*., 0 V) is applied to the EOM, the laser beam goes through a polarizing beamsplitter (PBS, PBS121, Thorlabs) and is magnified by a pair of relay lenses. When a high voltage (*i*.*e*., 260 V) is applied to the EOM, the laser beam is reflected by the PBS, and a pair of axicon lenses (AX2520-A, Thorlabs) is used to convert the collimated Gaussian beam into a ring-shaped beam for generating the Bessel beam. As a result, the EOM switches the beam profile between the Gaussian and Bessel beam pulse by pulse. The boxed inset in Fig. 2 illustrates the laser excitation scheme. By triggering the AOM and EOM at 10 kHz and 20 kHz, respectively, the pulse train emitted by the laser operating at a 40-kHz pulse repetition rate is packaged into multiple 4-pulse packets, each of which consists of one 532-nm Gaussian pulse, one 532-nm Bessel pulse, one 558-nm Gaussian pulse, and one 558-nm Bessel pulse. In the end, an objective lens (OL, AC254-050-A, Thorlabs) focuses all the beams onto the imaging object through a correction lens (CL, LA1207-A, Thorlabs) and the central opening of a ring-shaped ultrasonic transducer (UT, inner diameter: 1.1 mm; outer diameter: 3.0 mm; focal length: 4.4 mm; center frequency: 40 MHz; 6-dB bandwidth: 69%). The optically excited ultrasonic waves are detected by the ring transducer, amplified by a low-noise amplifier (HD28082, HD Communications), and acquired by a high-speed data acquisition board (DAQ, ATS9350, AlazarTech) at 500 MS/s. For raster scan, the object is mounted on a two-axis scanner, which consists of two motorized linear stages (L-509, Physik Instrumente). For acoustic coupling, the UT and CL are immersed in a water tank and a thin layer of ultrasound gel (Aquasonic CLEAR, Parker Laboratories) is applied between the object and a piece of transparent polyethylene membrane at the bottom of the water tank. A field-programmable gate array (PCIe-7842r, National Instruments) is programmed to synchronize the laser, AOM, EOM, motorized linear stage, and DAQ during the image acquisition.

## III. Results

First, to characterize the performance of the cGAN-powered Bessel-beam multi-parametric PAM, a phantom experiment was performed. Specifically, 3-μm polystyrene black dyed microspheres (24292-15, Polysciences) dissolved in deionized water (20%, v/v) were imaged over a depth range of 600 μm. Figure 3(a) shows the PAM images of the microspheres acquired with the Gaussian-beam and Bessel-beam excitation (before and after the cGAN-based processing), respectively. As shown in the top row, the images acquired with the Gaussian-beam excitation show a significant degradation in the resolution when the microspheres are out of focus, while the microsphere images acquired with the Bessel-beam excitation maintain a near-constant resolution and SNR across the entire depth range (middle row). Moreover, with the aid of cGAN-based post-processing, the artifacts induced by the side-lobes of Bessel beam are effectively removed (bottom row). The cross-sectional profiles of a representative microsphere imaged with the Gaussian- and Bessel-beam excitation (before and after the cGAN-based processing) are shown in Fig. 3(b), from which the in-plane resolutions under the three different settings (*i*.*e*., Gaussian-beam excitation, Bessel-beam excitation, and Bessel-beam excitation plus cGAN-based processing) are estimated to be 3.7, 4.5, and 4.0 μm, respectively, by quantifying the FWHM values of the cross-sectional profiles. In addition, 20 individual microspheres at each depth are selected to quantify the average FWHM values. Figure 3(c) shows the FWHM value as a function of the distance from the focal plane. In this phantom study, the cGAN-powered Bessel-beam PAM demonstrates an extended depth of focus (600 μm) with excellent image quality.

**Fig. 3.**
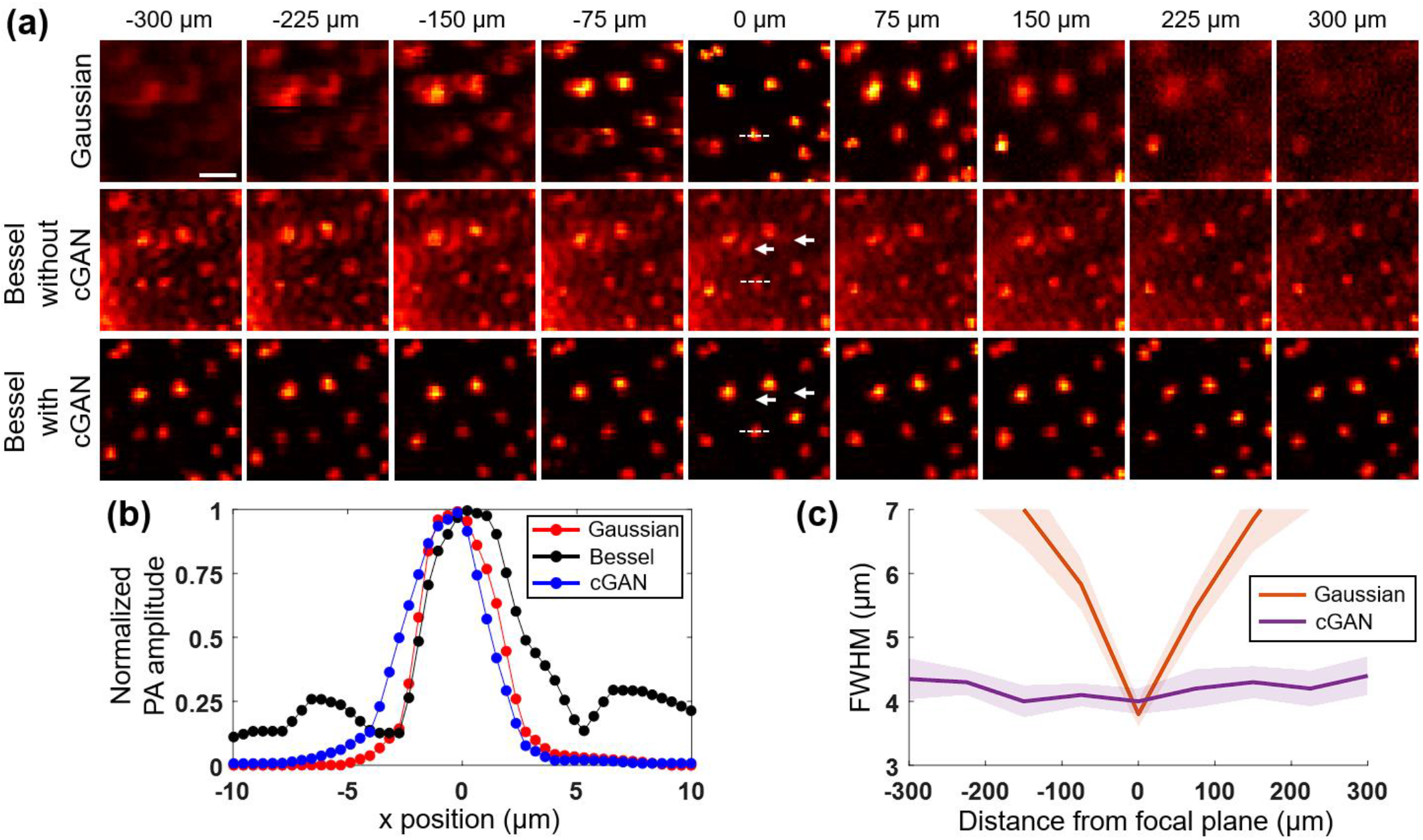
Characterization of the performance of the cGAN-powered Bessel-beam multi-parametric PAM. (a) PAM images of microspheres acquired with the Gaussian-beam and Bessel-beam excitation (before and after the cGAN-based image processing), respectively, over a depth range of -300 μm to 300 μm with respect to the focal plane (*i*.*e*., 0 μm). Scale bar: 20 µm. (b) Cross-sectional profiles of a representative microsphere imaged with the Gaussian beam and Bessel beam (before and after the cGAN-based processing), as indicated along the white dashed line in (a). The FWHM values of the cross-sectional profiles are 3.7, 4.5, and 4.0 μm for the Gaussian-beam excitation, Bessel-beam excitation, and Bessel-beam excitation plus cGAN-based post-processing, respectively. (c) FWHM value as a function of the distance from the focal plane. At each depth, 20 microspheres are selected for the quantification. The shading represents the standard deviation.

To examine whether the cGAN-powered Bessel-beam multi-parametric PAM can ensure accurate functional imaging over an extended depth range compared to the Gaussian-beam multi-parametric PAM, a side-by-side comparison and a cortex-wide recording were carried out *in vivo* in the mouse brain. Two male CD-1 mice (14 weeks old, Charles River Laboratories) were used for the experiments. Under general anesthesia (vaporized isoflurane: 2% for induction and 1.5% for maintenance), an open-skull window was created. Then, the ultrasound gel was applied between the cranial window and the membrane at the bottom of the water tank. The body temperature of the animal was maintained at 37°C by a heating pad (DCT-15, Kent Scientific). All experimental procedures were carried out in conformity with the animal protocol approved by the Institutional Animal Care and Use Committee at Washington University in St. Louis.

In Fig. 4(a), the top three rows are the multi-parametric images of C_Hb_, sO_2_, and CBF over a ∼2×2 mm^2^ region of interest acquired by the Gaussian-beam multi-parametric PAM at different depths, and the bottom three rows are corresponding images acquired by the Bessel-beam multi-parametric PAM and post-processed by the cGAN. It can be observed that the multi-parametric measurements made with the Gaussian-beam excitation quickly deviate from that in the focal plane, which is in contrast to the Bessel-beam case. The reason is that the validity of the multi-parametric measurements relies highly on the spatial resolution and SNR, as shown by our recent findings [8]. For the Gaussian beam, the optical spot spreads rapidly when out of focus, resulting in a fast-decaying SNR. On the other hand, the Bessel beam is non-diffractive. The focal spot size remains almost unchanged over a considerable range along the propagation direction. After the process of cGAN, the degradations of resolution and SNR are much slower. Fig 4(b) shows the percentage errors of the C_Hb_, sO_2_, and CBF measurements in 20 vessel segments as a function of depth for both systems (Gaussian *vs*. Bessel). For the sO_2_ measurement, the Bessel-beam multi-parametric PAM shows an error of <3.5% over an extended depth range of 600 μm, ∼13.3 times lower than the measurement error of the Gaussian-beam multi-parametric PAM. For the C_Hb_ and CBF measurements, even when the measurements were taken 300 μm away from the focal plane, the Bessel-beam multi-parametric PAM can still provide accurate readouts (percentage errors are ∼4.4% and ∼4.1% for CBF and C_Hb_, respectively), which are much more reliable than the Gaussian-beam system (percentage errors are ∼255% and ∼210% for CBF and C_Hb_, respectively).

**Fig. 4.**
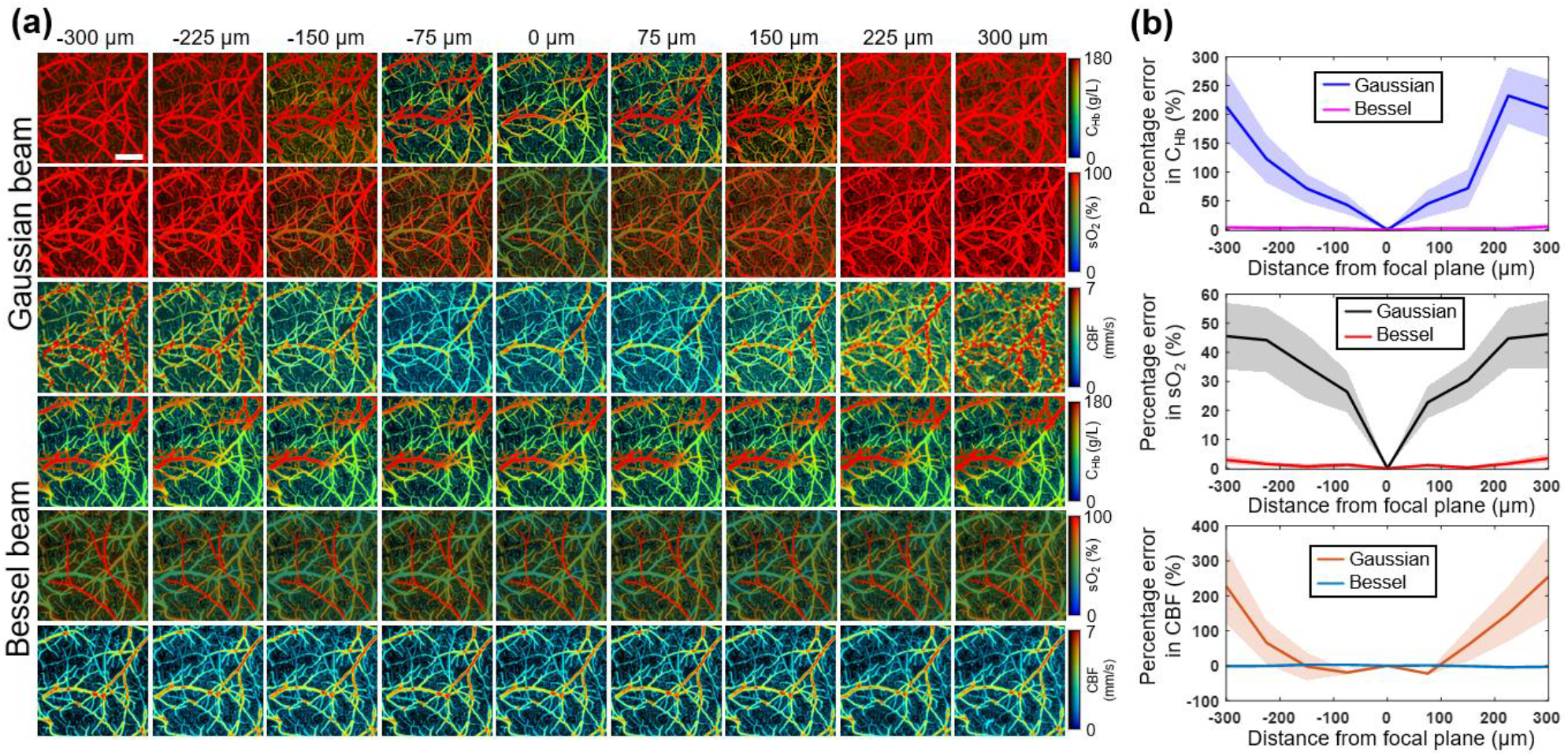
(a) Multi-parametric PAM images of C_Hb_, sO_2_, and CBF acquired with the Gaussian-beam and Bessel-beam excitation, respectively, over a depth range of -300 μm to 300 μm with respect to the focal plane. Scale bar: 500 µm. (b) Percentage errors in the measurements of C_Hb_, sO_2_, and CBF as a function of the distance from the focal plane. The shading represents the standard deviation.

Figure 5 shows cortex-wide imaging of the mouse brain by the Bessel and Gaussian-beam multi-parametric PAM. From left to right, the first column are the normalized PA amplitude and multi-parametric images of C_Hb_, sO_2_, and CBF over a ∼4×6 mm^2^ region of interest acquired by the Bessel-beam multi-parametric PAM. The close-ups of the lower right corner in the blue dashed box are shown right next to the column. The second column shows corresponding images acquired by the Gaussian-beam multi-parametric PAM. The last column on the right is the ground truth acquired by moving the Gaussian-beam to the focal plane. As expected, the Gaussian-beam multi-parametric PAM can get reasonable measurement in the center of the brain due to small deviation from the focal plane. However, on the two sides of the brain, the measurement becomes invalid as the deviation increases. On the other hand, the Bessel-beam multi-parametric PAM maintains a reliable quantification across the entire width of the mouse cortex, as confirmed by the ground truth. Such an extended depth of focus would enable many new biological and physiological studies, especially where imaging over a large field of view with an uneven surface is needed.

**Fig. 5.**
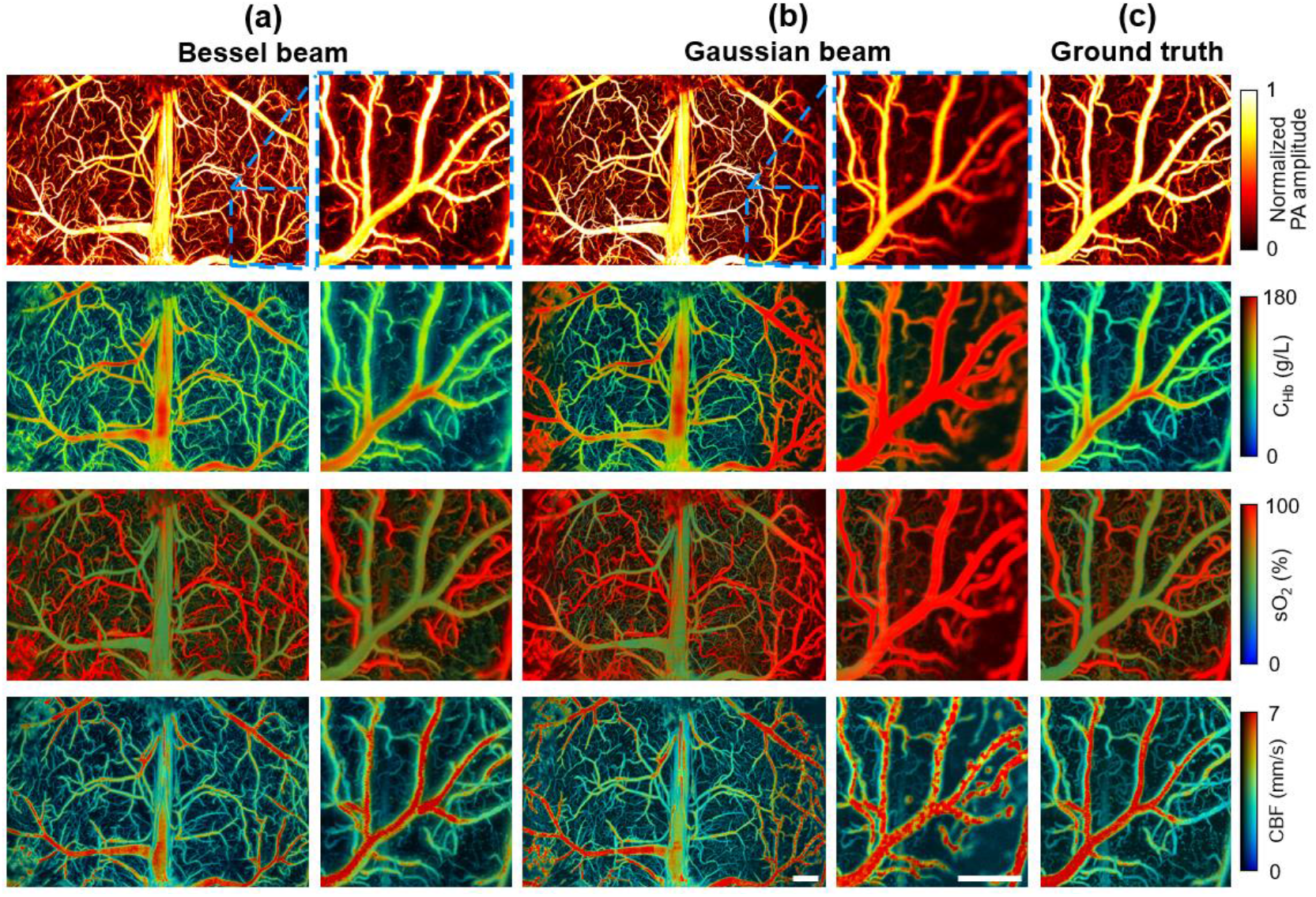
Cortex-wide multi-parametric PAM images of normalized photoacoustic amplitude, C_Hb_, sO_2_, and CBF acquired by (a) Bessel-beam and (b) Gaussian-beam excitation, respectively. The close-up of the blue boxed area is shown on the right-hand side. (c) Multi-parametric PAM images of the same region acquired with Gaussian beam in the focal plane. Scale bars: 500 μm.

## IV. Conclusion

In conclusion, we have developed a new multi-parametric PAM system based on Bessel-beam excitation and cGAN-based deep learning. Side-by-side comparison in the live mouse brain shows the advantages of the Bessel-beam excitation over the conventional Gaussian-beam excitation in maintaining accurate hemodynamic quantifications over the extended depth of focus. This new technique may find useful applications in large-field volumetric imaging of biological tissues with uneven surfaces, such as the brain and the tumor. Also, it can be integrated with the real-time contour scan we previously developed to further improve the tolerance of multi-parametric PAM for out-of-focus issues.

